# Genomic analysis of *P* elements in natural populations of *Drosophila melanogaster*

**DOI:** 10.1101/107169

**Authors:** Casey M. Bergman, Michael G. Nelson, Vladyslav Bondarenko, Iryna A. Kozeretska

## Abstract

The *Drosophila melanogaster P* transposable element provides one of the best cases of horizontal transfer of a mobile DNA sequence in eukaryotes. Invasion of natural populations by the *P* element has led to a syndrome of phenotypes known as P-M hybrid dysgenesis that emerges when strains differing in their *P* element composition mate and produce offspring. Despite extensive research on many aspects of *P* element biology, many questions remain about the genomic basis of variation in P-M dysgenesis phenotypes in natural populations. Here we compare gonadal dysgenesis phenotypes and genomic *P* element predictions for isofemale strains obtained from three worldwide populations of *D. melanogaster* to illuminate the molecular basis of natural variation in cytotype status. We show that the number of predicted *P* element insertions in genome sequences from isofemale strains is highly correlated across different bioinformatics methods, but the absolute number of insertions per strain is sensitive to method and filtering strategies. Regardless of method used, we find that the number of euchromatic *P* element insertions predicted per strain varies significantly across populations, with strains from a North American population having fewer *P* element insertions than strains from populations sampled in Europe or Africa. Despite these geographic differences, numbers of euchromatic *P* element insertions are not strongly correlated with the degree of gonadal dysgenesis exhibited by an isofemale strain. Thus, variation in *P* element insertion numbers across different populations does not necessarily lead to corresponding geographic differences in gonadal dysgenesis phenotypes. Additionally, we show that pool-seq samples can uncover population differences in the number of *P* element insertions observed from isofemale lines, but that efforts to rigorously detect differences in the number of *P* elements across populations using pool-seq data must properly control for read depth per strain. Our work supports the view that euchromatic *P* element copy number is not sufficient to explain variation in gonadal dysgenesis across strains of *D. melanogaster*, and informs future efforts to decode the genomic basis of geographic and temporal differences in *P* element induced phenotypes.

## Introduction

A substantial portion of eukaryotic genomes is represented by transposable elements (TEs). These TE families include those that colonized genomes long ago during the evolution of the host species and groups, but also those that have appeared in their host genomes recently. One of the best examples of a newly acquired TE is the *P* element in *Drosophila melanogaster* which is thought to have been acquired at least 70 years ago as a result of a horizontal transmission event from *D. willistoni* (Anxolabehere, Kidwell & Periquet, 1988; Daniels et al., 1990), a species that inhabits South America, the Caribbean, and southern parts of North America. Laboratory strains of *D. melanogaster* established from wild populations before the 1950s did not contain *P* element, while by the late 1970s this TE family was found in all natural populations worldwide (Anxolabehere, Kidwell & Periquet, 1988).

Classical work has shown that the presence of *P* elements induces a number of phenotypes in *D. melnaogaster* that can be characterized by the so-called “P-M hybrid dysgenesis” assay (Kidwell, Kidwell & Sved, 1977). Among the most prominent *P* element induced phenotypes is gonadal dysgenesis (GD), which is the key marker determining P-M status in particular strains of flies (Kidwell, Kidwell & Sved, 1977; Engels & Preston, 1980). In the P-M system, fly strains can be categorized as follows: P-strains have the ability to activate and repress *P* element transposition, P’-strains only have the ability to activate *P* element transposition, Q-strains only have the ability to repress *P* element transposition, and M-strains have neither the ability to activate or repress *P* element transposition (Kidwell, Kidwell & Sved, 1977; Engels & Preston, 1980; Quesneville & Anxolabéhère, 1998). M-strains that carry *P* element sequences in their genome are called M’-strains, while true M-strains are completely devoid of *P* elements (Bingham, Kidwell & Rubin, 1982). GD phenotypes were originally proposed to be mediated by repressor proteins encoded by full length *P* elements or truncated *P* elements that prevent *P* element transposition and subsequent DNA damage (Rio, 2002). Other work posits that these phenotypes mostly arise due to RNAi-based repression mediated by piRNAs produced by telomeric *P* elements and the effects are amplified by RNAs produced by other *P* elements (Simmons et al., 2014, 2015). More recently, some authors have questioned the classical view that GD phenotypes are caused solely by *P* elements or whether other factors may be involved (Zakharenko & Ignatenko, 2014; Ignatenko et al., 2015).

To better understand *P* element invasion dynamics and the molecular mechanisms that underlie P-M hybrid dysgenesis, many studies have surveyed variation in GD phenotypes across natural populations of *D. melanogaster* (Kidwell, Frydryk & Novy, 1983; Anxolabehere et al., 1985; Boussy & Kidwell, 1987; Anxolabehere, Kidwell & Periquet, 1988; Anxolabehere et al., 1988; Boussy et al., 1988; Gamo et al., 1990; Matsuura et al., 1993; Itoh et al., 1999; Bonnivard & Higuet, 1999; Itoh et al., 2001, 2004, 2007; Ogura et al., 2007; Onder & Bozcuk, 2012; Onder & Kasap, 2014; Ignatenko et al., 2015). These studies reveal that in most natural strains of *D. melanogaster* are P, Q, or M’, but that there can be substantial variation in the frequency of GD phenotypes within and between populations. In addition, variation among populations in GD phenotypes is thought to be relatively stable since their initial transitions from M cytotype to P, Q and M’ cytotypes (Gamo et al., 1990; Matsuura et al., 1993; Boussy et al., 1998; Bonnivard & Higuet, 1999; Itoh et al., 2001, 2004, 2007; Ogura et al., 2007). For example, Australian populations demonstrate a north-south cline of the frequency of various GD phenotypes (Boussy et al., 1987), which underwent only minor changes in the frequencies of truncated and full-size copies of the *P* element a decade later (Ogura et al., 2007).

A number of studies have also used Southern blotting, *in situ* hybridization to polytene chromosomes, or PCR to understand how the genomic composition of *P* elements varies qualitatively in relation to GD phenotypes (Todo et al., 1984; Engels, 1984; Boussy et al., 1988; Itoh et al., 1999, 2001; Itoh & Boussy, 2002; Ruiz & Carareto, 2003; Itoh et al., 2007; Onder & Kasap, 2014; Ignatenko et al., 2015). These studies have revealed that, irrespective of GD phenotype, the majority of *D. melanogaster* strains harbor multiple copies of full-length *P* elements (FP) along with multiple copies of the truncated repressor element known as “KP”, suggesting a complex relationship between the presence of different types of *P* elements in a genome and GD phenotypes. Attempts to quantify the relationship between absolute *P* element copy number or FP/KP ratios and GD phenotypes have revealed weak or no correlations between genomic *P* element composition and GD phenotypes (Todo et al., 1984; Engels, 1984; Boussy et al., 1988; Ronsseray, Lehmann & Anxolabéhère, 1989; Rasmusson et al., 1990; Itoh et al., 1999; Bonnivard & Higuet, 1999; Itoh & Boussy, 2002; Itoh et al., 2004, 2007). However, these conclusions rely on estimates of *P* element copy number based on low-resolution hybridization data.

The recent widespread availability of whole-genome shotgun sequences for *D. melanogaster* offers the possibility of new insights into the relationship between *P* element genomic content and GD phenotypes with unprecedented resolution. To date, hundreds of re-sequenced genomes of *D. melanogaster* exist and can be freely used for population and genomic analyses (Mackay et al., 2012; Pool et al., 2012; Lack et al., 2015; Bergman & Haddrill, 2015; Grenier et al., 2015; Lack et al., 2016). Moreover, a number of computational algorithms have been designed for *de novo* TE insertion discovery, annotation, and population analysis in *Drosophila* (Kofler, Betancourt & Schlötterer, 2012; Linheiro & Bergman, 2012; Cridland et al., 2013; Nakagome et al., 2014; Zhuang et al., 2014; Rahman et al., 2015). Comparison of different methods for detecting TEs in *Drosophila* NGS data has shown that they identify different subsets of TE insertions (Song et al., 2014; Rahman et al., 2015), and thus determining which TE detection method is best for specific biological applications remains an area of active research (Ewing, 2015; Rishishwar, Marino-Ramirez & Jordan, 2016).

To better understand the molecular basis of differences in cytotype status among populations, we investigated the relationship between GD phenotypes and *P* element predictions in whole genome shotgun sequences from three worldwide populations of *D. melanogaster*. By combining previously published GD assay data (Ignatenko et al., 2015) with *P* element predictions (this study) from genomic data of the same strains (Bergman & Haddrill, 2015), we show that the number of euchromatic *P* elements is not correlated with the degree of a GD phenotype exhibited by a strain. Furthermore, we show that populations can differ significantly in their euchromatic *P* element content, yet show similar distributions of GD phenotypes. We also investigate several bioinformatics strategies for detecting *P* element insertions in strain-specific and pooled genomic data to ensure robustness of our conclusions and help guide further genomic analysis. Our work supports previous conclusions that euchromatic *P* element copy number is not sufficient to explain variation in GD phenotypes, and informs future efforts to decode the genomic basis of differences in *P* element induced phenotypes over time and space.

## Materials and Methods

### Gonadal dysgenesis phenotypes

We re-analyzed GD assay data from (Ignatenko et al., 2015) for 43 isofemale strains of *D. melanogaster* from three geographic regions: North America (Athens, Georgia, USA), Europe (Montpellier, France), and Africa (Accra, Ghana), described in (Verspoor & Haddrill, 2011). Definitions of A and A* crosses in (Ignatenko et al., 2015) are inverted relative to those proposed by (Engels & Preston, 1980), and were standardized prior to re-analysis here. Cross A measures the activity of tester strain males mated to M-strain Canton-S females; cross A* measures the susceptibility of tester females mated to a P-strain Harwich males. P, P’, Q, and M-strains were defined according to (Kidwell, Frydryk & Novy, 1983; Quesneville & Anxolabéhère, 1998).

### Genome-wide identification of *P* element insertions

Whole genome shotgun sequences from the same *D. melanogaster* strains used for GD assays were downloaded from the European Nucleotide Archive (ERP009059) (Bergman & Haddrill, 2015). These genomic data were collected using a uniform library preparation and sequencing strategy (thus mitigating many possible technical artifacts) and include data for both individual isofemale strains and pools of single flies from isofemale strains (see (Bergman & Haddrill, 2015) for details). In total, 43 isofemale strain genomes from (Bergman & Haddrill, 2015) were analyzed that had GD data in (Ignatenko et al., 2015). Two pool-seq samples were analyzed for each population [N. America (15 and 30 strains), Europe (20 and 39 strains), and Africa (15 and 32 strains)]. Pool-seq samples contain one individual each from the same strains that have isofemale genomic data, plus additional strains that do not have GD data reported in (Ignatenko et al., 2015).

*P* element insertions were identified by TEMP (revision d2500b904e2020d6a1075347b398525ede5feae1; (Zhuang et al., 2014)) and RetroSeq (revision 700d4f76a3b996686652866f2b81fefc6f0241e0; (Keane, Wong & Adams, 2013)) using the McClintock pipeline (revision 3ef173049360d99aaf7d13233f9d663691c73935; (http://github.com/bergmanlab/mcclintock; Nelson, Linheiro & Bergman, 2016)). McClintock was run across the major chromosome arms (chr2L, chr2R, chr3L, chr3R, chr4, chrY, and chrX) of the UCSC dm6 version of the Release 6 reference genome (Hoskins et al., 2015) using the following options:-C -m “retroseq temp” -i -p 12 -b. Reference TE annotations needed for TEMP were generated automatically by McClintock using RepeatMasker (version open-4.0.6). The *D. melanogaster* TE library used by McClintock to predict reference and non-reference TE insertions is a slightly modified version of the Berkeley *Drosophila* Genome Project TE data set v9.4.1 (https://github.com/cbergman/transposons/blob/master/misc/D_mel_transposon_sequence_set.fa; described in (Sackton et al., 2009)).

In addition to providing “raw” output for each component method in the standardized zero-based BED6 format, McClintock generates “filtered” output tailored for each method (Nelson, Linheiro & Bergman, 2016). McClintock filters TEMP output to: (i) eliminate predictions where the start or end coordinates had negative values; (ii) retain predictions where there is sequence evidence supporting both ends of an insertion; and (iii) retain predictions that have a ratio of reads supporting the insertion to non-supporting reads of >1/10. Likewise, McClintock filters RetroSeq output to: (i) eliminate predictions where two different TE families shared the same coordinates; and (ii) retain predictions assigned a call status of greater than or equal to six as defined by (Keane, Wong & Adams, 2013).

Graphical and statistical analyses were performed in the R programming environment (version 3.3.2).

## Results

### Comparison of cytotype status and *P* element insertions in individual strains from North America, Europe and Africa

To address whether genomic data can be used to understand how cytotype status varies geographically and temporally, we identified *P* element insertions in publicly available genome sequences (Bergman & Haddrill, 2015) for a panel of 43 isofemale strains from three global regions with previously-published GD phenotypes (Ignatenko et al., 2015). As reported in (Ignatenko et al., 2015), isofemale strains from these populations were mainly P, M and Q (Figure 1, Table 1). Based on genomic analysis, all strains in these populations that are defined phenotypically as M are actually M’ (File S1). For N. American and African populations, the degree of activity tends to vary more across strains relative to susceptibility (Table 1, Figure S1A–B). However, we found no evidence for systematic differences across populations in the degree of activity (One-way ANOVA; *F*=0.06, 2 d.f., *P*=0.94) or susceptibility (One-way ANOVA; *F*=1.66, 2 d.f., *P*=0.2) (Figure S1A–B).

**Figure 1.**
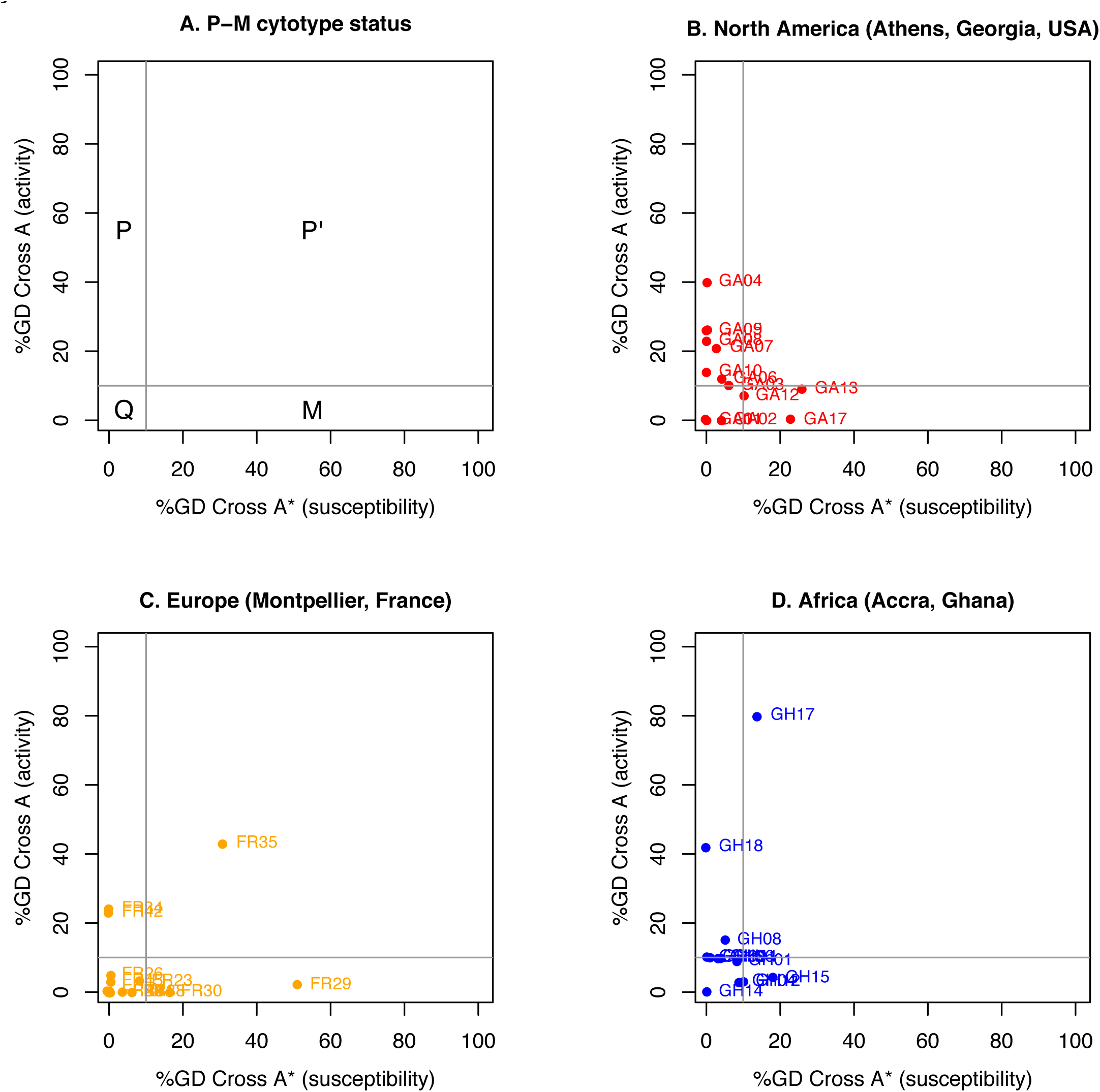
Results of GD tests for isofemale strains from natural populations from North America,Europe and Africa. %GD for cross A (tester strain males versus M-strain Canton-S females, vertical axis) and cross A* (P-strain Harwich males versus tester strain females, horizontal axis) are based on data reported in (Ignatenko et al., 2015). Cross A and A* labels in (Ignatenko et al., 2015) are inverted relative to those proposed by (Engels & Preston, 1980) and were converted to standard labels prior to analysis here. Each dot represents an isofemale strain. The P-M status for various sectors of GD phenotypic space defined by A and A* crosses are according to (Kidwell, Frydryk & Novy, 1983; Quesneville & Anxolabéhère, 1998) are shown in panel A.

**Table 1.** Gonadal dysgenesis (GD) levels and P-M status for isofemale strains of *D. melanogaster* obtained from natural populations in North America, Europe and Africa. %GD for cross A (tester strain males versus M-strain Canton-S females) and cross A* (P-strain Harwich males versus tester strain females) are based on data reported in (Ignatenko et al., 2015). Cross A and A* labels in (Ignatenko et al., 2015) are inverted relative to those proposed by (Engels & Preston, 1980) and were converted to standard labels prior to analysis here. P-M status for individual strains is according to (Ignatenko et al., 2015). Phenotypically M-strains are in fact M’-strains based on analysis of genomic data (File S1, File S2).

| Population | Year of<br>collection | Cross A<br>(%GD±SD) | Cross A*<br>(%GD±SD) | #<br>strains | M | Q | P' | P |
| --- | --- | --- | --- | --- | --- | --- | --- | --- |
| N. America | 2009 | 13.4±12.3 | 5.4±8.6 | 14 | 3 | 4 | 0 | 7 |
| Europe | 2010 | 6.1±12.2 | 6.8±14.0 | 17 | 2 | 12 | 1 | 2 |
| Africa | 2010 | 16.3±22.8 | 6.0±5.9 | 12 | 1 | 8 | 1 | 2 |
| Total |  |  |  | 43 | 6 | 24 | 2 | 11 |

We predicted *P* element insertions in the genomes of these isofemale strains using two independent bioinformatics methods – TEMP (Zhuang et al., 2014) and RetroSeq (Keane, Wong & Adams, 2013) – to ensure that our conclusions are not dependent on the idiosyncrasies of a single TE detection software package (File S1, File S2). We also investigated the effects of the default filtering of TEMP and RetroSeq output performed by McClintock (Nelson, Linheiro & Bergman, 2016), a meta-pipeline that runs and parses multiple TE insertion detection methods. We note that neither TEMP nor RetroSeq attempt to differentiate full-length from truncated insertions in their output, and we omitted heterochromatic contigs from our analysis. Overall numbers of euchromatic *P* elements predicted per strain by the different methods were well correlated across strains, regardless of the method of analysis and filtering (*r*≥0.712) (Figure 2). The highest correlation among methods was for the filtered TEMP and filtered RetroSeq datasets (*r*=0.945). McClintock filtering substantially reduced the average number of TEMP predictions for all three populations, bringing them more closely in line with the numbers predicted by RetroSeq (Table 2). These results suggest that the filtering steps performed by McClintock improve the consistency of TE predictions made by TEMP and RetroSeq on isofemale strains, and that the filtered data are more likely to reflect the true *P* element content of these lines.

**Figure 2.**
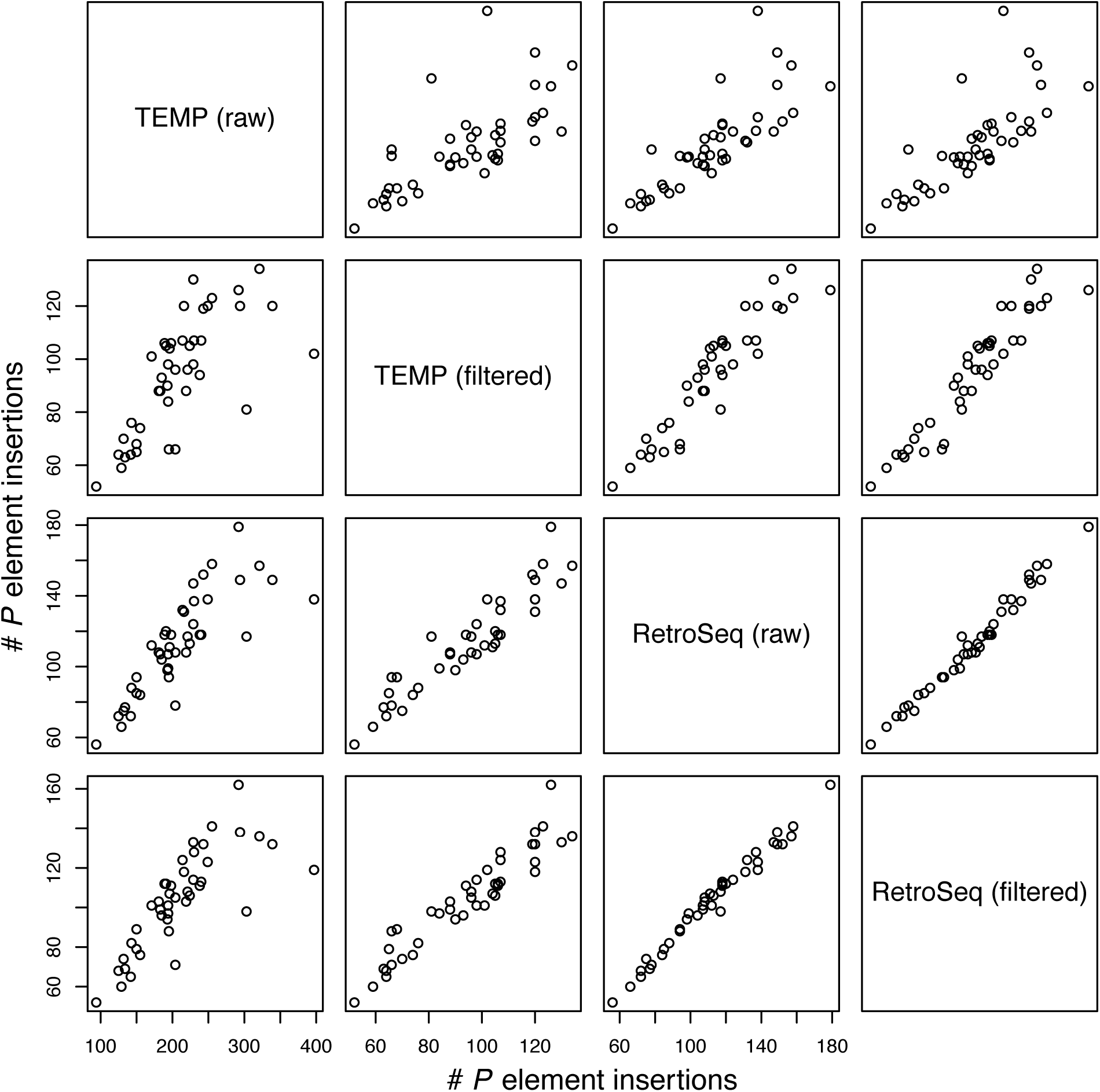
Correlation among methods in the numbers of predicted *P* element insertions for a worldwidesample of isofemale strains from North America, Europe and Africa. Numbers of *P* elements predicted by TEMP or RetroSeq shown are before (raw) and after (filtered) filtering by McClintock (see methods for details). Each circle represents an isofemale strain. Note that the scales on the x-axis and y-axis vary for each method.

**Table 2.** Average numbers (±S.D.) of *P*element insertions identified by TEMP and RetroSeq inisofemale strains from three worldwide populations of *D. melanogaster*. Columns labeled raw and filtered represent output generated by each method before or after default filtering by McClintock, respectively (see Materials and Methods for details).

| Population | # strains | TEMP<br>raw | TEMP<br>filtered | RetroSeq<br>raw | RetroSeq<br>filtered |
| --- | --- | --- | --- | --- | --- |
| N. America | 14 | 159.8±50.3 | 68.3±9.1 | 83.3±16.3 | 76.7±14.3 |
| Europe | 17 | 232.6±59.6 | 106.2±13.1 | 124.4±21.2 | 114.5±17.4 |
| Africa | 12 | 232.8±41.2 | 106.8±14.5 | 129.3±20.0 | 119.2±14.5 |

The average number of *P* element insertions predicted per strain for all three populations is shown in Table 2. In general, McClintock filtered predictions data suggests these isofemale lines contain ~70-120 *P* element insertion sites, which is roughly 2-fold higher than the 30-50 copies per haploid genome estimated from Southern blotting (Bingham, Kidwell & Rubin, 1982; Ronsseray, Lehmann & Anxolabéhère, 1989; Bonnivard & Higuet, 1999; Itoh & Boussy, 2002). These results are consistent with increased resolution of *P* element predictions based on genomic data plus residual heterozygosity due to incomplete inbreeding in these strains (Lack et al., 2016). In contrast to the lack of population difference observed at the phenotypic level, genomic data shows clear differences in the numbers of euchromatic *P* element insertions in strains from North American populations relative to the European and African populations, regardless of the TE detection method and filtering (One-way ANOVA; *F*>9.26; 2 d.f., *P*<5e-4) (Table 2; Figure S1C–F; Figure 3). Taken together with the GD data, these results suggest that population-level differences in the abundance of *P* elements per strain do not necessarily lead to population-level differences in the frequency of GD phenotypes.

Integrating published GD data with our genomic predictions at the level of individual strains, we directly tested whether the number of *P* elements per strain is associated with either GD phenotype. We found that neither the degree of activity (cross A) nor the degree of susceptibility (cross A*) was significantly linearly correlated with the filtered number of predictions made by TEMP or RetroSeq (p>0.11; Figure 3). Similar results were obtained using the raw output of these methods as well (Figure S2). These results confirm results at the population level above and suggest that there is no simple relationship between the total number of euchromatic *P* elements and GD phenotypes at the level of individual strains.

**Figure 3.**
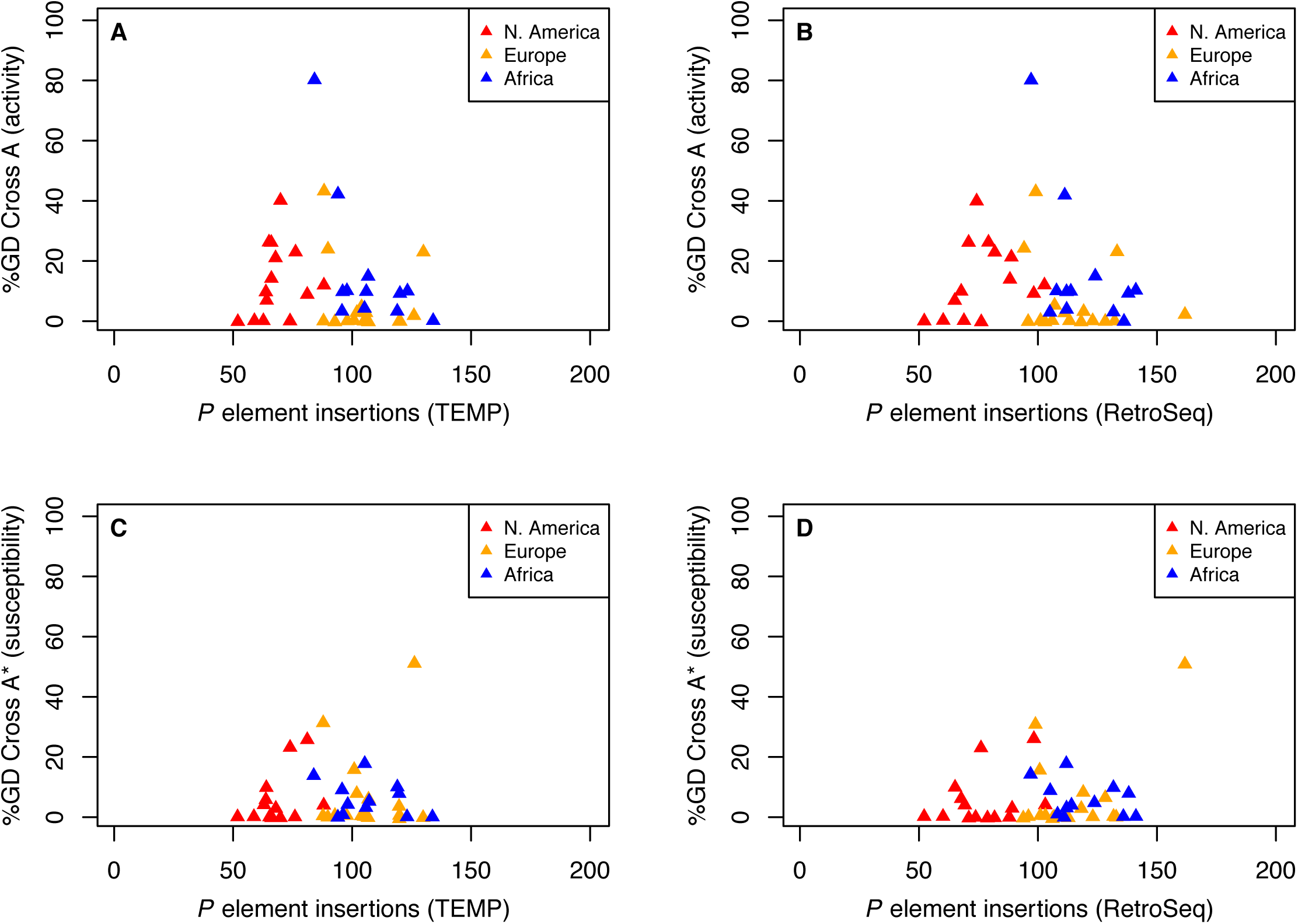
Relationship between %GD in A and A* crosses and filtered numbers of euchromatic *P* element insertions identified by TEMP or RetroSeq for isofemale strains from natural populations from North America, Europe and Africa. %GD data are from [46] and use the same standardized definitions as in Figure 1. Numbers of *P* elements predicted by TEMP or RetroSeq shown here are after filtering by McClintock (see Materials and Methods for details). Analogous results for unfiltered raw output of TEMP or RetroSeq are shown in Figure S2. Each triangle represents an isofemale strain.

### Population difference in *P* element insertion numbers can be observed in pool-seq samples

Pooled-strain sequencing (pool-seq) is a cost-effective strategy to sample genomic variation across large numbers of strains and populations (Schlotterer et al., 2014). To address whether the differences among populations we observed in the number of *P* elements predicted in isofemale strain data are also seen in pool-seq data, we predicted *P* element insertions in pool-seq samples from the same populations (Table 3, File S2). Two pool-seq samples are available for each population that differ in the number of individuals (one per isofemale strain) used: North America (n=15 and n=30), Europe (n=20 and n=39), and Africa (n=15 and n=32). The smaller pools from each population include one individual from the same isofemale strains analyzed above; the larger pools contain one individual from the same strains as the smaller pools, plus individuals from additional isofemale strains from the same population that were not sequenced as isofemale strains. Thus, the smaller pool-seq samples are a nested subset of the larger pool-seq samples, and pool-seq samples from the same population are not fully independent from one another.

**Table 3.** Numbers of *P* element insertions identified by TEMP and RetroSeq in pool-seq samples from three worldwide populations of *D. melanogaster*. Numbers in parentheses are numbers of insertions scaled by the number of strains in the pool. Columns labeled raw and filtered represent output generated by each method before or after default filtering by McClintock, respectively (see Materials and Methods for details).

| Population | # strains | TEMP<br>raw | TEMP<br>filtered | RetroSeq<br>raw | RetroSeq<br>filtered |
| --- | --- | --- | --- | --- | --- |
| N. America | 15 | 684 (45.6) | 53 (3.5) | 312 (20.8) | 219 (14.6) |
| N. America | 30 | 1,101 (36.7) | 27 (0.9) | 259 (8.6) | 159 (5.3) |
| Europe | 20 | 1,003 (50.1) | 85 (4.2) | 372 (18.6) | 245 (12.2) |
| Europe | 39 | 1,395 (35.8) | 46 (1.2) | 278 (7.1) | 171 (4.4) |
| Africa | 15 | 958 (63.9) | 110 (7.3) | 505 (33.7) | 348 (23.2) |
| Africa | 32 | 1,681 (52.5) | 28 (0.9) | 329 (10.3) | 193 (6.0) |

The numbers of *P* element insertions identified by TEMP and RetroSeq in pool-seq samples are given for all three populations in Table 3. In the raw output, TEMP predicted more insertions in larger strain pools relative to smaller strain pools, as expected for a method designed to capture TE insertions that are polymorphic within a sample (Zhuang et al., 2014). However, McClintock-filtered TEMP output generated between 9-fold and 60-fold fewer insertions per sample in the pool-seq output relative to raw output, as well as fewer insertions overall in the larger strain pools relative to the smaller strain pools. These effects are likely because of the McClintock requirement for TEMP predictions to have the proportion of reads supporting the insertion to non-supporting reads to be >10%. In contrast, McClintock filtering reduced the total number of RetroSeq predictions by less than 2-fold, and fewer insertion sites were predicted for the larger strain pools in both raw and filtered RetroSeq output.

When the number of strains in a pool-seq sample was used as a scaling factor, pool-seq samples yield many fewer *P* element predictions per strain than the average average number of *P* elements predicted from the same isofemale strains (Table 2, Table 3). This is expected because of lower sequencing per strain depth in the pooled samples relative to the isofemale line samples. Similarly, the scaled data shows fewer insertions were predicted per strain in the larger pools relative to smaller pools for all populations regardless of method or filtering (Table 3). This effect arises because larger and smaller strain pool samples contain similar numbers of reads (~44 million read pairs per sample), and thus larger strain pools have fewer reads per strain. Because pooled samples contain the same strains as isofemale lines and because smaller pools contain a subset of the same strains that are present in larger pools, these results suggest a dilution effect for *P* element detection in pool-seq samples: at a fixed sequencing coverage, *P* element insertions that are predicted in samples with higher coverage cannot be detected in samples with lower coverage, even though they are in fact present in the sample.

In spite of this dilution effect, African pool-seq samples tend to have more insertions per strain than North American samples (Table 3), similar to what is seen in the isofemale strain datasets (Figure 3, Table 2). This result is most clearly demonstrated for the comparison between North American and African samples which each had 15 strains, where the African sample has more predicted insertions regardless of TE detection method and filtering. These results suggest that, if dilution effects are properly controlled for, pool-seq samples can capture general trends among populations in total *P* element insertion numbers that are seen in isofemale strain sequencing.

## Discussion

Here we performed a detailed analysis of *P* element content in genomes of isofemale strains and pool-seq samples from three worldwide populations of *D. melanogaster* with published GD phenotypes. Our results allowed us to draw several conclusions about the detection of *P* element insertions in *D. melanogaster* population genomic data and the genomic basis of GD phenotypes that can be used to inform future studies.

For samples derived from isofemale strains, we find that two different TE detection methods (TEMP and RetroSeq) generate well-correlated numbers of *P* element predictions per strain (Figure 2), but that filtering by McClintock improves the overall correlation between these methods (mainly by reducing the number of presumably false positive TEMP predictions). In contrast, analysis of pool-seq samples revealed larger differences between TE detection methods and a larger effect of McClintock filtering (primarily because of how insertions that are polymorphic within a sample are handled). Pool-seq samples yield fewer predicted insertions per strain than the average number of insertions per strain for the same set of isofemale strains, most likely because of the lower per-strain sequencing coverage in pool-seq samples. Similarly, we found that there is a diminishing return on the number of *P* element insertions detected per strain in pool-seq samples for a given sequencing coverage, regardless of method or filtering. These dilution effects mean it will be difficult to compare *P* element predictions from pool-seq data with those from isofemale strains or to compare pool-seq samples to each other, unless the read depth per strain in the pool is carefully controlled. We note that the observation of diminishing returns for a fixed level of coverage in pool-seq samples does not contradict previous claims that (with increasing total sequencing coverage) there appears to be no diminishing return on detection of new TE insertions in *D. melanogaster* pool-seq samples (Rahman et al., 2015).

Regardless of prediction method, we found no simple linear relationship between the strength of GD phenotypes and the number of euchromatic *P* element insertions across isofemale strains (Figure 3). Our results are consistent with previous attempts to connect total numbers of *P* elements in a genome to GD phenotypes, which found weak or no correlations using Southern blotting or *in situ* hybridization to polytene chromosomes (Todo et al., 1984; Engels, 1984; Boussy et al., 1988; Ronsseray, Lehmann & Anxolabéhère, 1989; Rasmusson et al., 1990; Itoh et al., 1999; Bonnivard & Higuet, 1999; Itoh & Boussy, 2002; Itoh et al., 2004, 2007). Assuming that GD assays using single reference strains provide robust insight into the GD phenotypes of these natural strains, our results are at face value consistent with recent arguments that the *P* element may not be the primary determinant of hybrid dysgenesis (Zakharenko & Ignatenko, 2014). However, our results are also consistent with GD phenotypes being determined by one or more active full-length *P* element insertions found in specific locations in the euchromatin, or by the relative abundance of full-length and truncated repressor elements, rather than overall copy numbers (which includes both active and inactive copies). Alternatively, the lack of correlation between the number of *P* element insertions and GD phenotypes may result from noise in the data due to the genomic sequence data not being of sufficient depth in these samples, or the GD assays having substantial experimental variation across lines.

Nevertheless, we did observe differences among populations in the number of predicted *P* element insertions per strain (Figure 3), even though no strong differences were observed in the levels of GD phenotypes across these populations. Specifically, strains from the North American population had the fewest predicted *P* element insertions, regardless of the TE detection method or filtering (Figure 3, Figure S1). This result is somewhat unexpected given that the *P* element is thought to have first been horizontally transferred into a North American population before invading the rest of the world (Anxolabehere, Kidwell & Periquet, 1988). This observation suggests that N. American populations may have evolved some form of copy number control not present in other populations. Evidence for fewer *P* element insertions per strain in the North American population could also be detected in pool-seq samples, especially when the number of strains per pool was controlled for (Table 3), indicating that pool-seq is a viable strategy for surveying differences in *P* element copy number across populations. Overall, our results show that it is possible to detect clear differences in euchromatic *P* element insertion profiles among populations using either isofemale strain or pool-seq genomic data, however interpreting how *P* element insertion site profiles relate to GD phenotypes at the strain or population level remains an open challenge.

## Acknowledgements

We thank Lyudmila Zakharenko, Justin Blumenstiel, and Nelson Lau for helpful comments on a previous version of the manuscript. This work was supported by Wellcome Trust PhD Studentship (096602/B/11/Z) to MGN, a Human Frontier Science Program Young Investigator grant RGY0093/2012 to CMB, a Biotechnology and Biological Sciences Research Council grant BB/L002817/1 (CMB), and free private repositories from GitHub (CMB).

## Supplemental Files

**File S1.** Tab separated value (TSV) formatted file with %GD data from A and A* crosses, P-M cytotype status, population, and numbers of predicted *P* elements in raw and filtered output from TEMP and RetroSeq, respectively, for 43 isofemale strains from three global regions. GD data are taken from (Ignatenko et al., 2015) and were standardized to definitions proposed by (Engels & Preston, 1980) prior to re-analysis here.

**File S2.** Zip archive of browser extensible data (BED) files of predicted *P* element locations in genome sequences from 50 isofemale strains and 6 pool-seq samples from three global regions. Each sample has four BED files corresponding to raw (*raw.bed) and filtered (*nonredundant.bed) output from TEMP and RetroSeq, respectively. BED files for 7 isofemale strains from (Bergman & Haddrill, 2015) are included here that do not have GD data in (Ignatenko et al., 2015) but are included in the pool-seq samples, allowing comparisons to be made between isofemale strains and pool-seq samples for the same set of strains.

**Figure S1.**
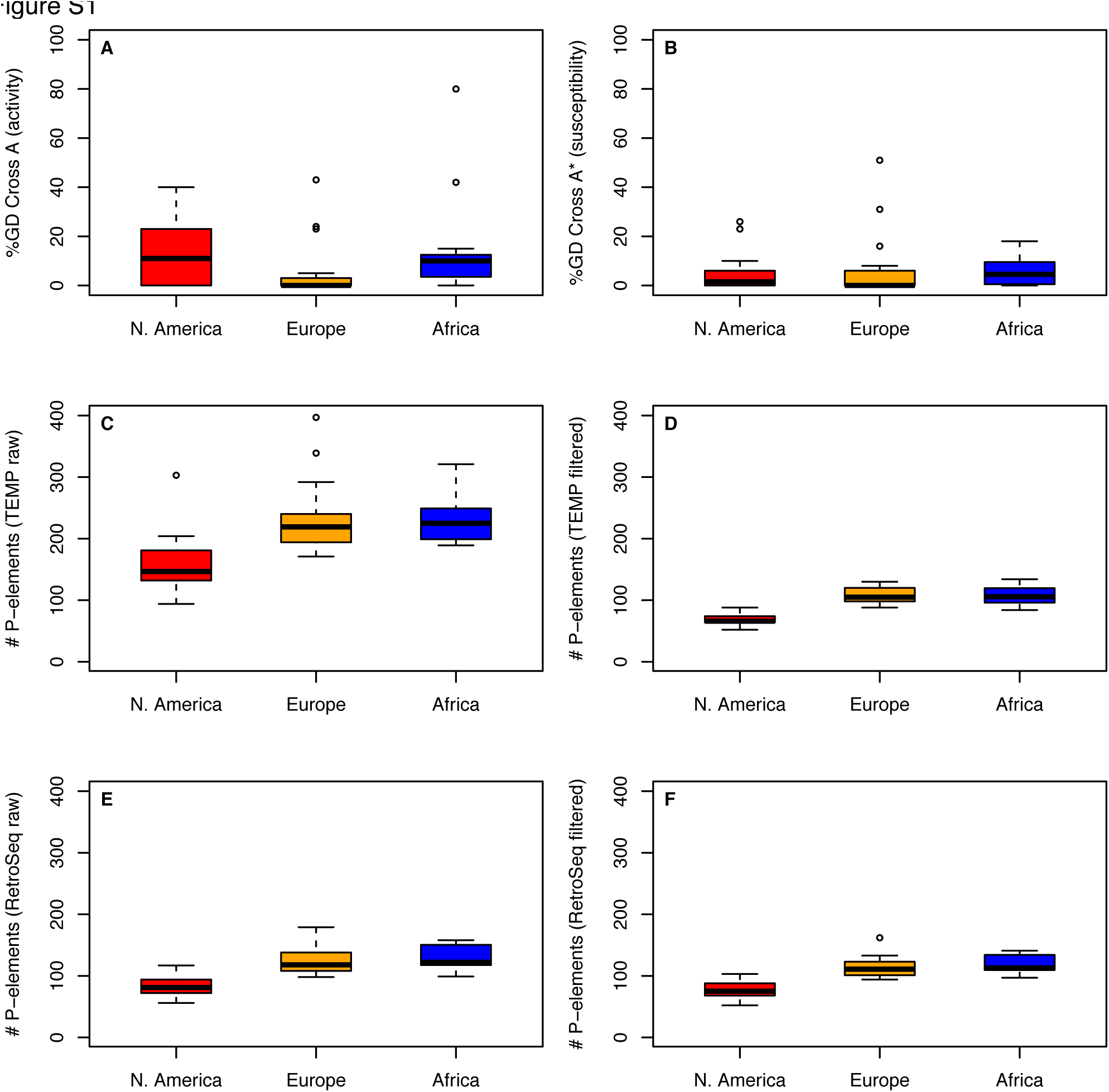
Distributions of %GD in A and A* crosses and numbers of predicted *P* element insertionsfor isofemale strains within and between populations from three worldwide regions. Distributions are shown as boxplots with black lines representing median values, boxes representing the interquartile range (IQR), whiskers representing the limits of values for strains that lie within 1.5 x IQR of the upper or lower quartiles, and circles representing strains that lie outside 1.5 x IQR of the upper or lower quartiles. GD data are taken from (Ignatenko et al., 2015) and were standardized to definitions proposed by (Engels & Preston, 1980) prior to re-analysis here. Numbers of *P* elements predicted by TEMP or RetroSeq shown are before (raw) and after (filtered) standard filtering by McClintock.

**Figure S2.**
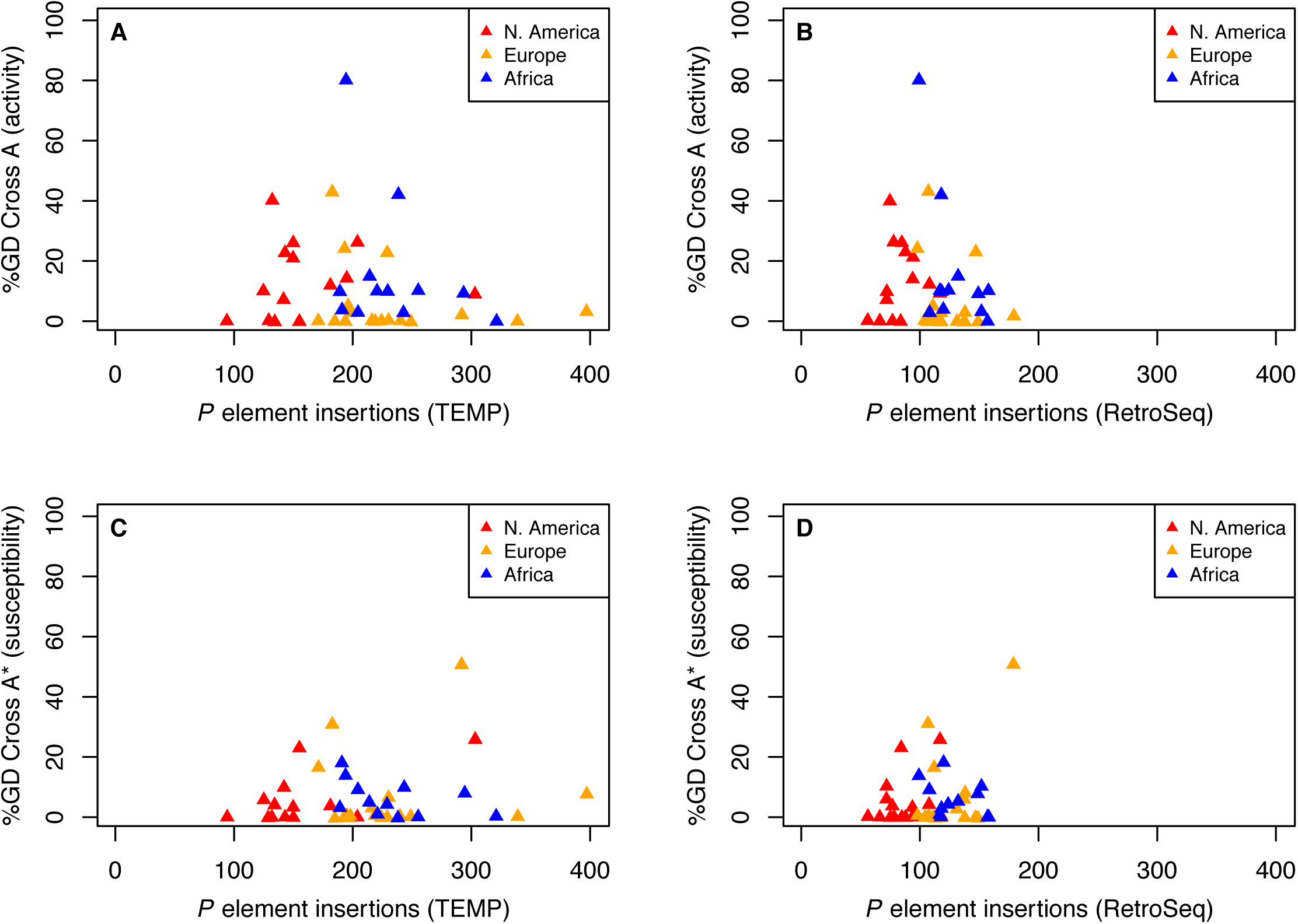
Relationship between %GD in A and A* crosses and raw numbers of euchromatic *P* element insertions identified by TEMP or RetroSeq for isofemale strains from natural populations from North America, Europe and Africa. %GD data are from (Ignatenko et al., 2015) and are the same standardized values as in Figure 1. Numbers of *P* elements predicted by TEMP or RetroSeq shown are raw output prior to standard filtering by McClintock. Analogous results for McClintock-filtered output of TEMP and RetroSeq are shown in Figure 3. Each triangle represents an isofemale strain.

